# A Modified Injector and Sample Acquisition Protocol Can Improve Data Quality and Reduce Inter-Instrument Variability of the Helios Mass Cytometer

**DOI:** 10.1101/600130

**Authors:** Brian H. Lee, Geoffrey Kelly, Shermineh Bradford, Melanie Davila, Xinzheng V. Guo, El-ad David Amir, Emily M. Thrash, Michael D. Solga, Joanne Lannigan, Brian Sellers, Julian Candia, John Tsang, Ruth R. Montgomery, Stanley J. Tamaki, Tara K. Sigdel, Minnie M. Sarwal, Lewis L. Lanier, Yuan Tian, Cheryl Kim, Denise Hinz, Bjoern Peters, Alessandro Sette, Adeeb H. Rahman

## Abstract

Mass cytometry is a powerful tool for high dimensional single cell characterization. Since the introduction of the first commercial CyTOF mass cytometer by DVS Sciences in 2009, mass cytometry technology has matured and become more widely utilized, with sequential platform upgrades designed to address specific limitations and to expand the capabilities of the platform. Fluidigm’s 3rd-generation Helios mass cytometer introduced a number of upgrades over the previous CyTOF2. One of these new features is a modified narrow bore sample injector that generates smaller ion clouds, which is expected to improve sensitivity and throughput. However, following rigorous testing we find that the narrow-bore sample injector may have unintended negative consequences on data quality and result in lower median signal intensities and higher coefficients of variation in antibody expression. We describe an alternative Helios acquisition protocol using a wider bore injector, which largely mitigates these data quality issues. We directly compare these two protocols in a multi-site study of 10 Helios instruments across 7 institutions and show that the modified protocol improves data quality and reduces inter-instrument variability. These findings highlight and address an important source of technical variability in mass cytometry experiments that is of particular relevance in the setting of multi-center studies.

## Introduction

Mass cytometry leverages time-of-flight inductively coupled plasma mass spectrometry to perform high-throughput high parameter single cell analysis. By substituting fluorochromes with metal isotopes, mass cytometry overcomes the limitations of fluorescence spectral overlap and allows the simultaneous analysis of over 40 different parameters on a single cell. This allows for deeper phenotypic and functional characterization of complex biological samples and has contributed to biological insights across a wide range of research areas.

Since the introduction of the first commercial CyTOF mass cytometer by DVS Sciences in 2009 (1), mass cytometry technology has matured and become more widely utilized, with sequential platform upgrades designed to address specific limitations and expand the capabilities of the platform. Fluidigm’s 3^rd^-generation Helios mass cytometer introduced a number of upgrades over the previous CyTOF2. These include a pneumatic sample introduction system that replaced the CyTOF2’s loop-based sample introduction system, which allowed for more consistent and efficient acquisition of large sample volumes. Another new feature of the Helios was a modification of the sample injector, a component that transports nebulized sample droplets from the spray chamber into the inductively couple plasma (ICP). The Helios injector has a narrower internal diameter than the CyTOF2 injector, resulting in the generation of smaller ion clouds, which, in principle offers two major advantages: 1) smaller ion clouds are less likely to fuse while transiting from the ICP to the ion optics, thus reducing the likelihood of ion-cloud doublet events at a given sample introduction rate; and 2) condensing a comparable number of cell-derived ions into a smaller cloud allows more efficient sampling of a cell’s content into the ion optics, and also allows each cloud to be measured in fewer pushes, thus increasing the relative signal per push and improving instrument sensitivity.

While the modified Helios injector offers some notable theoretical advantages over the CyTOF2, when we empirically tested these advantages by comparing the performance of the CyTOF2 and Helios platforms we noted discrepant behavior between synthetic beads used for instrument calibration and biological samples, and we found evidence of reduced overall data quality when analyzing cells using the standard Helios instrument configuration and acquisition protocol. These findings suggested that the narrow-bore injector may have unintended negative consequences to data quality. We further describe and characterize an alternative acquisition protocol using a recently-introduced modified wide bore (WB) injector in conjunction with a proprietary ionic cell acquisition solution (CAS) developed by Fluidigm, which largely mitigates these data quality issues. We directly compare the conventional NB and alternative WB Helios acquisition protocols in a multi-site study of 10 different instruments across 7 different institutions and show that the modified protocol improves data quality and reduces inter-instrument variability. Together, our findings highlight an important factor that may impact the quality of data acquired on many Helios cytometers under the standard instrument configuration and demonstrate the improvements that can be achieved using a modified acquisition protocol.

## Methods

### Samples and processing

PBMC samples were prepared from de-identified leukapheresis products obtained from the New York Blood Center. All protocols and procedures employed were reviewed and approved by the Mt. sinai institutional review committee. For the initial comparisons of the CyTOF2 and Helios cytometers and initial evaluations of the different Helios acquisition protocols, PBMCs were divided into multiple aliquots and stained with CD45 antibodies conjugated to unique metal tags across the detectable mass range together with an immune profiling panel to allow identification of major immune subsets. All antibodies were either purchased pre-conjugated from Fluidigm or conjugated in-house using commercial X8 polymer conjugation kits purchased from Fluidigm (supplementary table 1). After staining, samples were fixed with freshly-diluted 2.4% formaldehyde in PBS containing 0.02% saponin and iridium intercalator to label nucleated cells. In one experiment, samples were alternatively fixed with 0.05% glutaraldehyde or a combination of 2.4% formaldehyde and 0.05% glutaraldehyde in PBS (supplemental figure 2). Samples were acquired immediately after staining, or frozen in FBS containing 10% DMSO and stored at −80°C, which we and others have previously shown to be an effective strategy for long term storage of samples prior to CyTOF analysis (2,3). For the multi-site study, PBMCs from 5 independent donors were each barcoded with CD45 antibodies and pooled prior to staining with a standard immune profiling panel. The samples were then fixed with freshly-diluted 2.4% formaldehyde and stained with iridium intercalator to label nucleated cells. The stained and fixed cells were then divided into multiple aliquots and frozen as above. Frozen aliquots from the same batch of samples were distributed to 7 institutions and acquired on 10 separate mass cytometers using both acquisition protocols.

**Figure 1.**
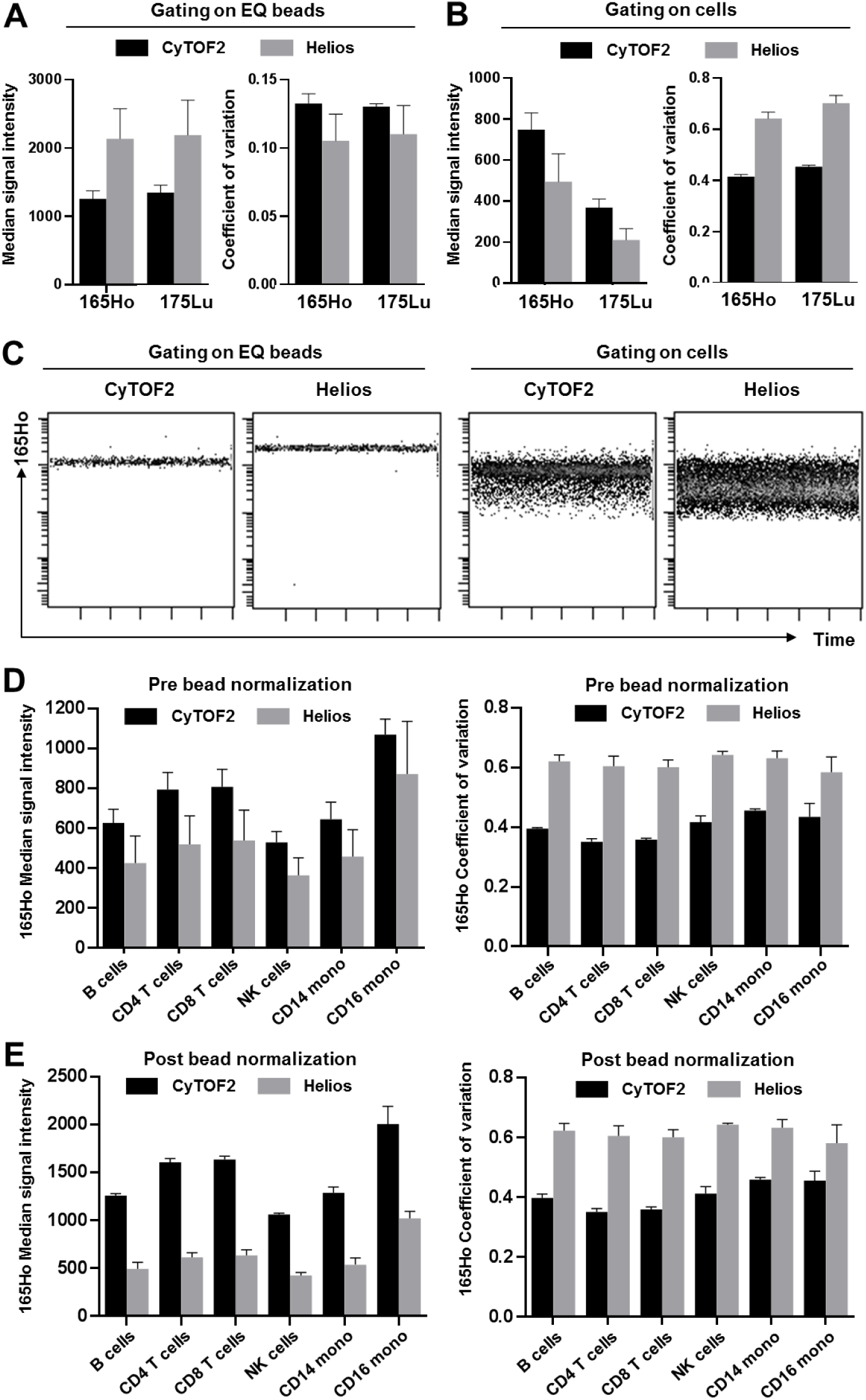
The Helios mass cytometer shows divergent performance when analyzing EQ beads and cells in the same sample. Replicate aliquots of CD45-barcoded PBMCs were spiked with EQ beads and run in parallel on a CyTOF2 and a Helios mass cytometer. (A) When gating specifically on EQ beads in the sample, the Helios shows higher median intensity and lower CVs for bead-associated channels including Ho165 and Lu175. (B) In contrast to the beads, expression of CD45-Ho165 and CD45-175Lu antibodies on cells in the same sample shows lower median intensity and higher CVs. (C) This higher CV does not reflect a progressive change in marker expression over time; higher variability persists consistently throughout the duration of the acquisition. (D) CD45 expression across defined gated cell subsets, highlighting that the reduced median and increased CV are not population-specific phenomenon. (E) Due to divergent behavior between cells and beads, the reduced signal intensity cannot be corrected by bead-based normalization, which also has no effect on marker CVs. Bars for all graphs represent the mean and S.D. of three aliquots of the same stained PBMC sample acquired independently on three separate days on one CyTOF2 and one Helios mass cytometer.

**Figure 2.**
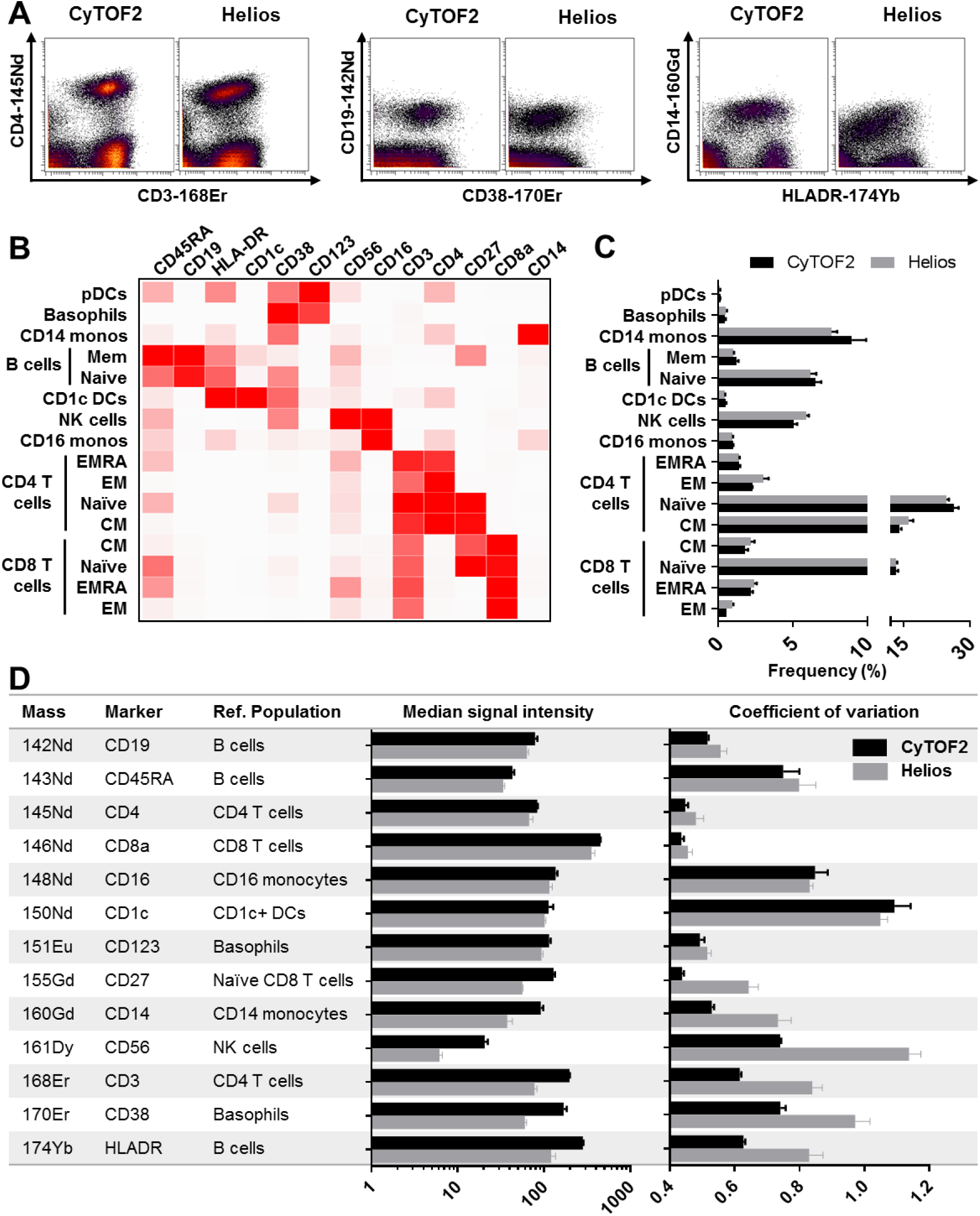
The Helios mass cytometer shows overall reduced cell staining quality relative to the CyTOF2. Replicate aliquots of the same PBMC sample were run in parallel on a CyTOF2 and a Helios mass cytometer. (A) Visualization of marker staining quality on traditional biaxial plots shows reduced intensity, higher CVs and overall poorer resolution of several markers on the Helios relative to the CyTOF2. (B-D) Data were clustered using SPADE and major populations were annotated and exported based on canonical marker expression patterns. (B) Heatmap of median marker expression across populations (scaled per marker). (C) Frequency of major immune populations measured using both injectors. (D) A specific positive reference population was defined for each marker in the panel and used to evaluate relative median marker intensity and population marker CV using both injectors. Bars for all graphs represent the mean and S.D. of three aliquots of the same stained PBMC sample acquired independently on three separate days on one CyTOF2 and one Helios mass cytometer.

### Data acquisition

PBMC samples were either acquired within 48hrs of staining or were frozen and stored at −80°C until thawing at RT immediately prior to acquisition. For the standard Helios acquisition protocol, the samples were washed twice with PBS + 0.2% BSA, once with MilliQ H_2_O, and resuspended at a concentration of 750,000 cell per ml (or as indicated in the figure legends) in MiliQ H_2_O containing a 1/10 dilution of EQ beads (Fluidigm). These samples were acquired using a standardized acquisition template following routine tuning and instrument optimization using the narrow bore Helios injector. For the modified acquisition protocol, the samples were washed twice with PBS + 0.2% BSA, once with CAS and resuspended at a concentration of 750,000 cell per ml in CAS containing a 1/10 dilution of EQ beads (Fluidigm). These samples were acquired using a standardized acquisition template following routine tuning and optimization using the new wide-bore Helios injector after equilibrating the instrument by running CAS for at least 5 minutes.

### Data analysis

FCS Files were normalized and concatenated as necessary using the Fluidigm acquisition software. In the case of multi-site studies, inconsistent channel naming was corrected using either *cytofcore* or *cytutils*. Files were uploaded to Cytobank(4) for manual bead exclusion, CD45-barcode deconvolution using Boolean gating, and manual gating analyses. Debarcoded samples were further analyzed by a combination of manual gating, SPADE clustering in Cytobank(5), or semi-automated flowSOM(6) clustering and cluster annotation using Astrolabe Diagnostics, a cloud-based cytometry analysis platform. Cell subset frequencies and the median signal intensities and CVs for each marker across each cell population were exported for subsequent analyses. Heatmap visualizations of median marker intensity across defined populations were performed by importing the median intensities of the SPADE-gated populations into Clustergrammer(7). To calculate changes in marker staining quality under the different protocols, a representative cell population was selected that was positive for each marker and the median marker intensity and marker CV for that population were exported for secondary statistical analyses using Prism (GraphPad Software). Marker-specific performance was determined by calculating the pairwise fold change in marker medians and CVs for each sample-population-marker combination (WB relative to NB). tSNE analyses were performed in Cytobank(8) and Jensen-Shannon Divergence between tSNE plots was calculated using *cytutils* (https://github.com/ismms-himc/cytutils).

## Results and Discussion

### Direct comparison of the CyTOF2 and Helios mass cytometers reveals divergent performance when evaluating reference beads and cell samples and indicates reduced cellular data quality on the Helios

Samples acquired by mass cytometry are typically spiked with a dilution of EQ beads (Fluidigm), which are synthetic beads containing known amounts of 5 metal isotopes that span the mass range of the instrument. These beads serve as a valuable resource to monitor instrument performance and normalize data to account for variations in intra- and inter-instrument performance(9). A key assumption in bead-based normalization is that the behavior of beads in a given sample approximates that of the cells in the same sample. When evaluating the performance of a new Helios mass cytometer compared to a CyTOF2 mass cytometer based on EQ beads, we noted that the Helios resulted in higher median signal intensities and lower coefficients of variance (CVs) (Fig. 1A), as predicted by the smaller ion clouds generated by the Helios injector, which was also apparent based on the reduced event length. However, when measuring the expression of cell-associated CD45 antibodies conjugated to the same isotopes as those used for the bead evaluations (CD45-165Ho and CD45-175Lu), we observed reduced median signal intensities and higher CVs (Fig. 1B) in the samples acquired on the Helios. Evaluating signal intensity over time indicated that the reduced cell-associated marker signal intensity and higher CV was apparent at the start of the Helios sample acquisition and remained stable over time, suggesting that it was not due to a progressive degradation of the sample over the acquisition period (Fig. 1C). This reduction in CD45 staining quality was also observed across multiple immune cells types suggesting that it was not a cell-type specific phenomenon (Fig. 1D). As noted previously, bead-based normalization can help to correct for fluctuations in instrument performance, but this assumes that cells and beads are similarly affected by these fluctuations. Given the discrepancy in signal performance between EQ beads and cells, bead-based normalization was not able to correct for the difference in data quality between the two instruments, having no effect on marker CVs and in fact further accentuating the differences between cell staining intensity between the two instruments (Fig. 1E).

This reduction in data quality was also apparent when evaluating markers to define immune cell subsets using conventional biaxial data visualizations (Fig. 2A). To more comprehensively characterize and quantify the impact on immune population identification and individual marker expression, the data were clustered using SPADE(5) and major populations were annotated based on canonical marker expression patterns and exported for further analysis (Fig. 2B). The relative frequencies of major cell lineages defined by highly distinct marker expression profiles were similar; however, when analyzing sub-clusters defined by more subtle differences in marker expression, the differences in data quality resulted in greater differences in reported frequencies (Fig. 2C). To quantify the impact on individual marker expression, we defined a specific population cluster positive for each marker in the panel and used this population as a reference to quantify the relative median signal intensity (Fig. 2D) and CV (Fig. 2E) of marker expression. This approach highlighted the relative reduction in marker intensity and corresponding increase in marker CV across a range of markers, with a more notable impact on markers at the higher end of the measured mass range.

### Characteristics of a new wide bore injector on the Helios system

At the 7th Annual Mass Cytometry Summit in May 2018, Fluidigm officially announced the release of a new “wide bore” (WB) injector and a proprietary “cell acquisition solution” (CAS) with a higher ionic content than water. The combination of the WB injector and the CAS buffer were proposed to improve data quality and stability on the Helios mass cytometer. The WB injector has an intermediate diameter between the original “narrow bore” (NB) Helios injector and the previous CyTOF2 injector (Fig. 3A). The use of this WB injector also requires a higher make-up gas flow than the NB injector (Fig. 3B), and results in larger ion clouds, as evidenced by average event lengths that are higher than those on the conventional Helios NB configuration but still smaller than those on the CyTOF2 (Fig. 3C). Larger ion clouds would be predicted to increase the likelihood of ion cloud fusion events, resulting in a higher rate of ion-cloud doublets. To evaluate this possibility, we utilized samples comprised of an equal ratio of cells barcoded with CD45 antibodies conjugated to two distinct isotopes: 169Tm and 175Lu, thereby allowing the identification of known 169+175+ doublets (Fig. 3D). When examining the Gaussian parameters that describe the ion cloud features associated with a given event, the known 169+175+ doublets exhibited higher Residual and Offset than the 169+175- and 175+169- populations, as would be expected of an ion cloud fusion event (Fig. 3E). It is important to note however, that there was considerable overlap in the relative Residual/Offset distributions between these three populations, suggesting that Gaussian parameters alone may not be sufficient to accurately identify all the doublets in a given sample. In addition to the defined 169+175+ doublets, we expected that the sample would also contain 169+169+ doublets, and 175+175+ doublets that would not be readily identifiable in the sample. However, the measured frequency of known doublets could be used to estimate frequency of total doublets. Assuming 4 doublet categories (169+169+; 169+175+; 175+169+; 175+175+), the frequency of total doublets = 2X the frequency of measured 169+175+ doublets. We confirmed the accuracy of this calculation by examining the relative proportions of other known cell-cell doublets (i.e., B cell-T cell doublets, T cell-monocyte doublets and B cell-monocyte doublets) and confirming that in each case, half of the overall doublets were found within the defined 169+175+ doublet population (Fig. 3E). We proceeded to serially dilute this barcoded sample across a range of concentrations and determined the average event rate/second at each concentration when acquired on the same Helios instrument tuned using the WB and NB injectors. We found that the WB injector resulted in a slightly lower average event rate for a given cell concentration, suggesting a lower acquisition efficiency than the NB injector (Fig. 3F). We further used the measured 169+175+ doublet rate and calculated the total inferred doublet frequency at a given event rate per second (Fig. 3G). These results confirmed that the larger ion clouds associated with the WB injector do result in higher doublet rates, and correspondingly suggest that an acquisition rate of ~250 events per second using the WB injector results in a doublet rate of ~9%, which is comparable to the doublet rate when running the same sample at ~400 events per second using the NB injector.

**Figure 3.**
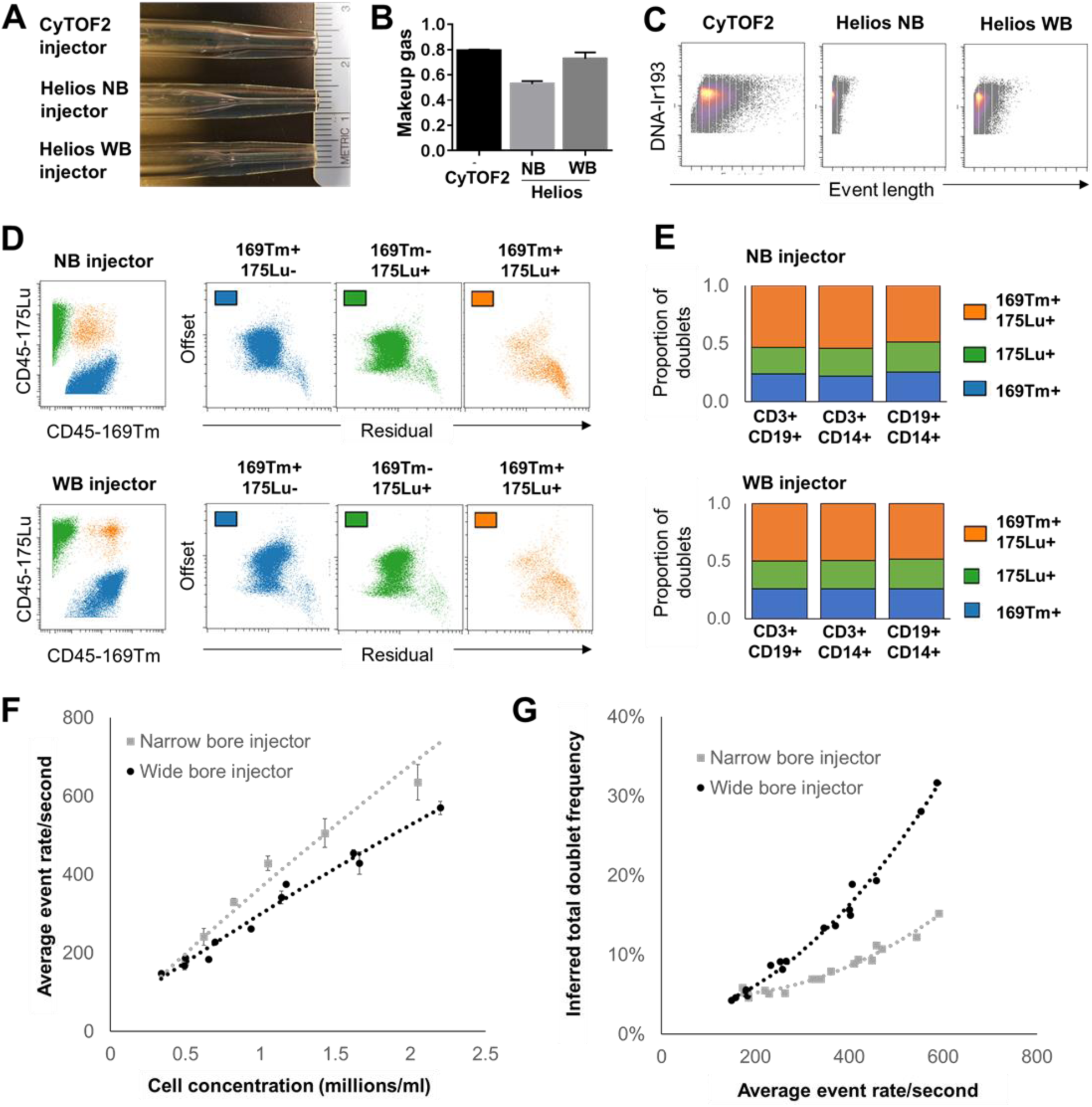
Acquisition characteristic of the narrow and wide bore injector Helios configurations. (A) Photographs of the CyTOF2, Helios NB and Helios WB injectors showing the intermediate internal bore diameter of the WB injector. (B) Average optimal makeup gas (L/min) with each of the injectors as determined by tuning with a fixed nebulizer gas rate of 0.17 L/min. (C) Representative event lengths for parallel sample aliquots acquired with each of the injectors. (D-G) Two aliquots of the same PBMC sample were barcoded with 169Tm and 175Lu-conjugated CD45-antibodies, which were combined at an equal ratio and acquired in parallel on the same Helios instrument using either the WB or NB injector. (D) 169Tm+175+ events were identified as known cross sample doublets and were confirmed to exhibit higher residual and offset gaussian parameter values. (E) The relative proportion of CD3+CD19+ (T cell-B cell), CD3+CD14+ (T cell-monocyte) and CD19+CD14+ (B cell-monocyte) doublets within the 169Tm+175Lu+ doublet population confirmed that the total doublet frequency could be accurately estimated as 2X the measured frequency of CD169+175+ doublets. (F) Average event rate for samples acquired at a given cell concentration using each injector. (G) Calculated total doublet frequency in relation to event acquisition rate using each injector. Data represent values from 3 replicate acquisitions of the same cell suspension diluted at each cell concentration and acquired on a single Helios instrument.

### A modified acquisition protocol using the WB injector improves data quality collected on the Helios

We next evaluated staining quality of parallel aliquots of the same sample acquired on the same Helios instrument using the original NB injector and WB injector using both H_2_O and CAS buffer during acquisition. Evaluation of marker intensity on biaxial plots confirmed our prior observations of reduced marker intensity and higher CV of the standard Helios relative to the CyTOF2, both of which were improved by the use of the WB injector and further improved when the WB injector was used in conjunction with CAS (Fig. 4A). Notably, the use of CAS was not recommended with the NB injector due to the greater likelihood of depositions clogging the narrow injector tip (Fluidigm, personal communication). Using a similar approach to that described in Fig. 2, we evaluated the staining quality of all the marker in the panel and found that the WB+CAS protocol restored Helios marker medians and CVs to levels comparable to or superior to those measured using the CyTOF2 (Fig. 4B). The relative improvement in data quality using the WB+CAS protocol was independently confirmed on a different Helios mass cytometer at a second site using PBMC samples from three donors, each of which was stained and acquired in triplicate (supplemental Fig. 1).

**Figure 4.**
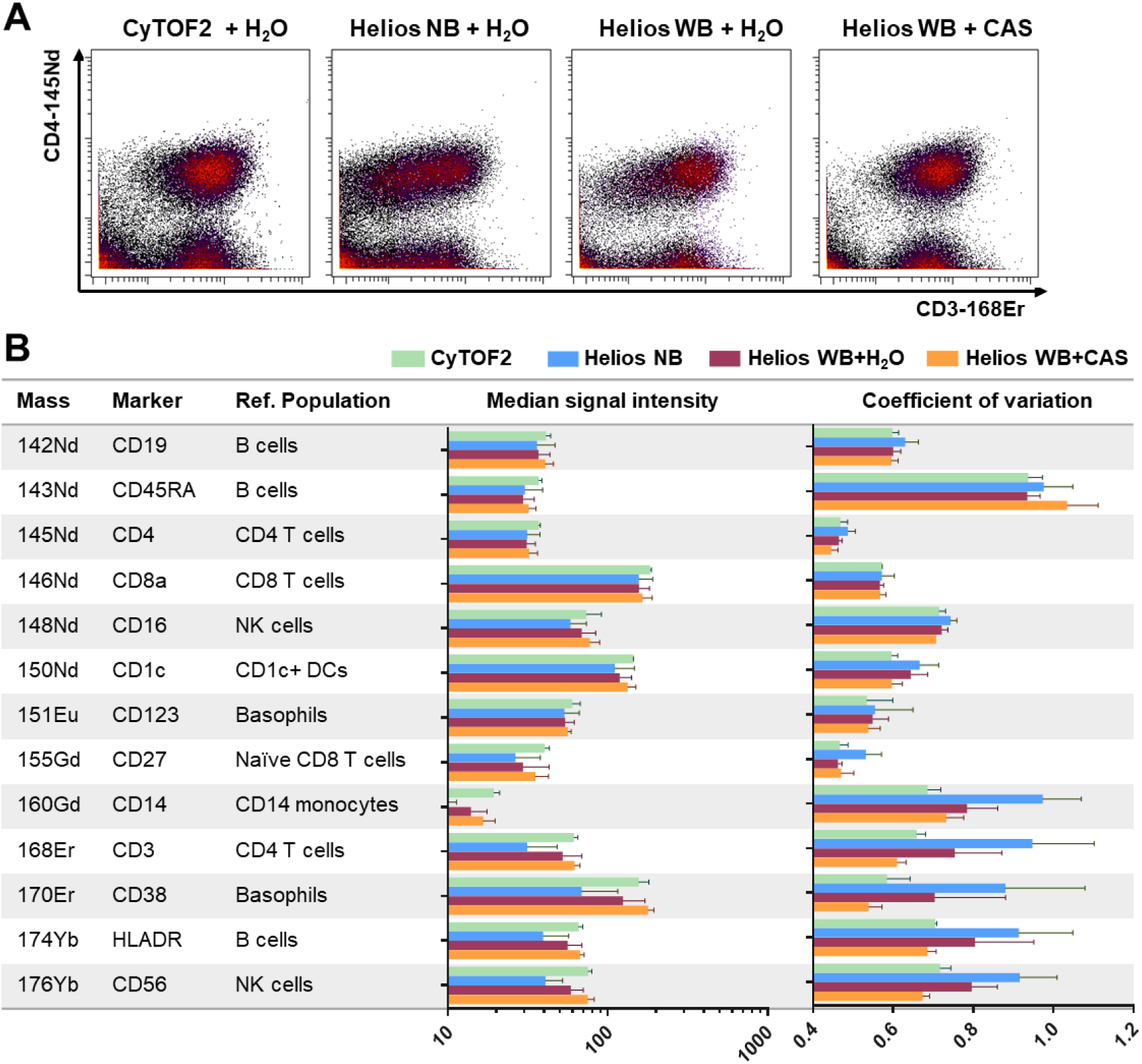
The WB injector + CAS protocol improves Helios data quality to levels comparable to the CyTOF2. Parallel aliquots of a CD14-depleted PBMC sample were stained and acquired on CyTOF2 or Helios mass cytometers using either the NB injector or WB injector with H_2_O or the WB injector with CAS. (A) Biaxial plots show relative improvement of CD3-168Er staining quality on the same Helios when using the WB injector, particularly in conjunction with CAS. (B) Evaluation of median marker intensity and marker CV for the indicated reference cell populations for each of the markers in the panel. Bars represent the mean and S.D. of three replicate samples run independently on two CyTOF2 instruments, one new Helios, and two CyTOF2s after upgrade to Helios specifications.

To address the potential that this phenomenon results from degradation of inadequately fixed samples, we prepared additional aliquots of a stained PBMC sample, which were fixed using either freshly-diluted 2.4% formaldehyde, 0.05% glutaraldehyde or a combination of 2.4% formaldehyde and 0.05% glutaraldehyde. The WB+CAS protocol resulted in overall improved data quality relative to the NB+H_2_O protocol for all samples across all three fixation conditions (supplementary Fig. 2). The WB+CAS protocol also improved data quality for samples that were acquired either immediately or after staining or following post-staining cryopreservation and storage in 10% DMSO in FBS. However, on some instruments we noted that sample cryopreservation further exacerbated the data quality artefacts associated with the NB injector, and that in these cases the WB+CAS protocol consequently resulted in greater relative improvement in data quality (supplementary Fig. 2). Overall, these findings suggest that while the data quality artefacts associated with the NB injector depend on both sample- and instrument-specific factors, the WB+CAS protocol has the potential to be broadly applicable to improve data quality for samples processed using a range of staining, fixation and storage protocols.

### A modified acquisition protocol reduces inter-instrument variability and improves data consistent in multi-site mass cytometry studies

To further validate these findings across a wider range of instruments we prepared a large batch of cells comprised of a pool of CD45-barcoded PBMCs from 5 independent donors stained with a general immunophenotyping panel. Aliquots of these stained cells were distributed to 7 institutions and acquired on a total of 10 Helios instruments using both the conventional NB injector + H_2_O protocol and the WB injector + CAS protocols, and the data were compiled for a central analysis. We found that the quality of data acquired using the NB + H_2_O protocol was highly variable across instruments, whereas the WB + CAS protocol resulted in more similar data across instruments (Fig. 5A). Consequently, while the WB + CAS showed an overall trend towards improvement in both population-specific marker intensity and marker CV across most instruments, this effect was also instrument specific, with some instruments showing a much larger change in performance than others (Fig. 5B). In this case of instruments that did experience a change in performance, this was generally reflected by an increase in median marker intensity, and a corresponding reduction in population specific marker CV (Fig. 5C). Given the considerable variation in the relative improvement in performance seen on individual instruments, we evaluated the tuning and performance parameters on each instrument to identify instrument parameters potentially associated with the improved performance (supplementary table 2). We observed a positive correlation between the overall improvement in staining quality and the Tb(159) dual counts following tuning with either the NB (Fig. 5D) or WB injectors (Fig. 5E). This suggests that Helios instruments showing the highest sensitivity are most likely to suffer from data quality artifacts with the NB injector, and are potentially most likely to benefit from adopting the WB + CAS protocol.

**Figure 5.**
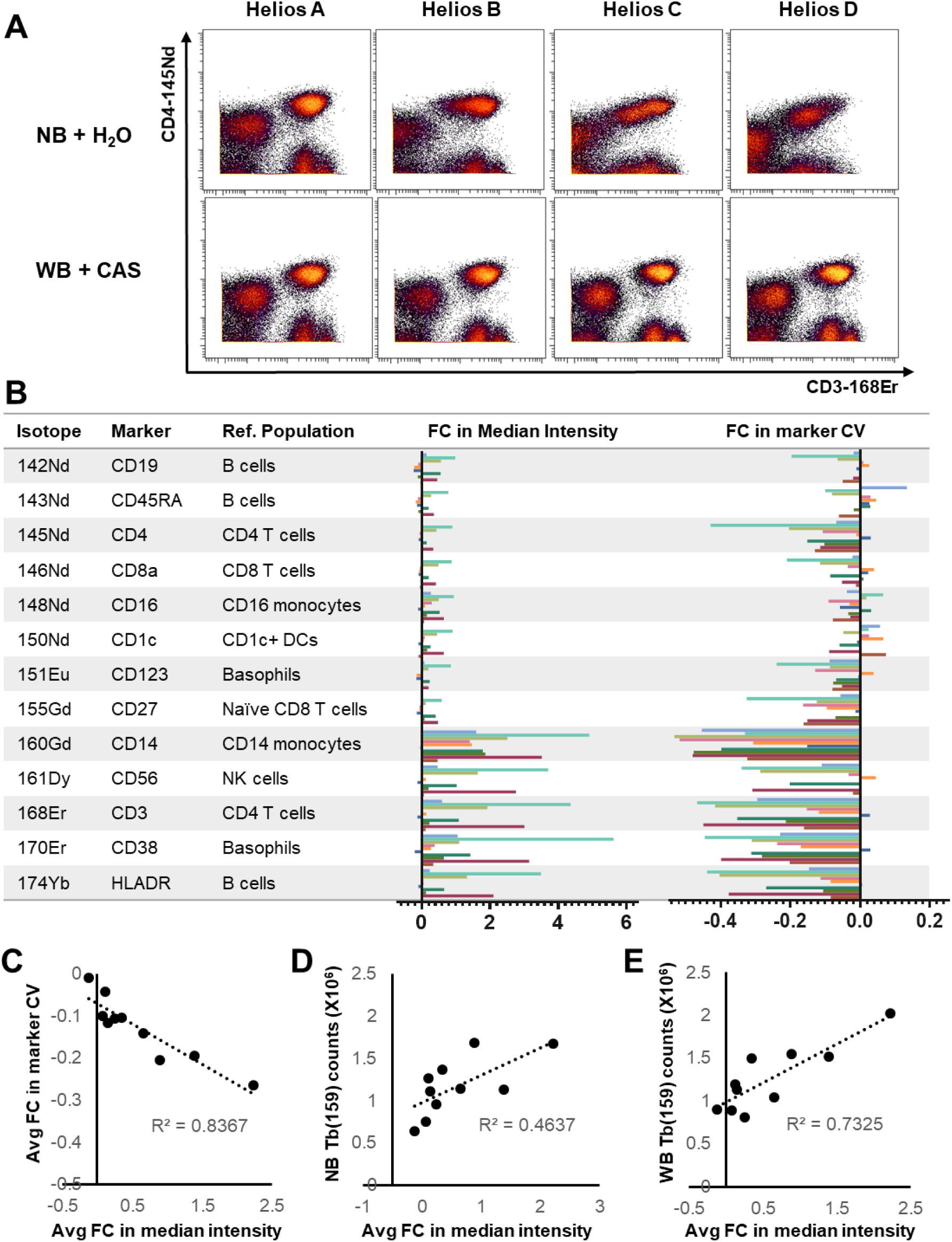
A multisite evaluation highlights that the relative impact of the different injector configurations is highly instrument dependent. Replicate aliquots of a pre-stained sample composed of CD45-barcded PBMCs from 5 healthy donors were distributed to 7 institutions and acquired on 10 independent Helios instruments using either the NB+H_2_O or the WB+CAS acquisition protocols. (A) Biaxial plots showing data from 4 representative instruments showing the variability in relative staining quality using each acquisition protocol across different instruments. (B) Calculation of the fold change in population-specific marker median and CV when transitioning from the NB+H_2_O to the WB+CAS protocol across all 10 instruments. Each bar represents an individual instrument. (C-E) Correlation between the overall fold change in median intensity (averaged across all markers) and overall change in marker CV (C), the Tb(159) dual counts measured after tuning with the NB injector (D) and the WB injector. Each point represents an individual instrument.

While the magnitude of the relative improvement was somewhat instrument specific, the overall result across all instruments was higher data consistency and reduced inter-instrument variability when using the WB + CAS acquisition protocol, as previously noted in Fig. 5A. This was apparent when evaluating sample-specific CD45 barcode staining quality in each of the five debarcoded PBMC samples, where the WB + CAS protocol resulted in more consistent median staining intensity (Fig. 6A) across all instruments. To evaluate the effect for the immunophenotyping markers in the panel, we performed tSNE analyses for each PBMC sample using all the markers in the panel (Fig. 6B) and calculated the pairwise Jensen-Shanon (JS) divergence to evaluate the relative difference between all plots for the NB+H_2_O and WB+CAS data respectively (Fig. 6C). The WB+CAS protocol resulted in significantly lower average JS divergence for each donor sample, indicating a reduction in overall inter-sample variability with the WB protocol. To further evaluate the impact on immune population identification we clustered and annotated the data from each sample using SPADE. While both protocols resulted in largely comparable population frequencies of major immune cell types (Fig. 6E), the WB+CAS protocol resulted in lower inter-instrument variation in the reported population frequencies (Fig. 6E). Similarly, the WB+CAS protocol also reduced inter-instrument variation in population-specific median marker intensity (Fig. 6F). Thus, the overall result is that the WB+CAS protocol reduces inter-instrument variation and results in more comparable data across instruments. This finding was further reproduced in a second multi-site study using an independent PBMC sample that was stained using an independent antibody cocktail and acquired on a subset of 6 of the same Helios instruments in the primary multi-site comparison, which again showed a reduction in inter-instrument variability using the WB+CAS protocol (supplemental figure 3).

**Figure 6.**
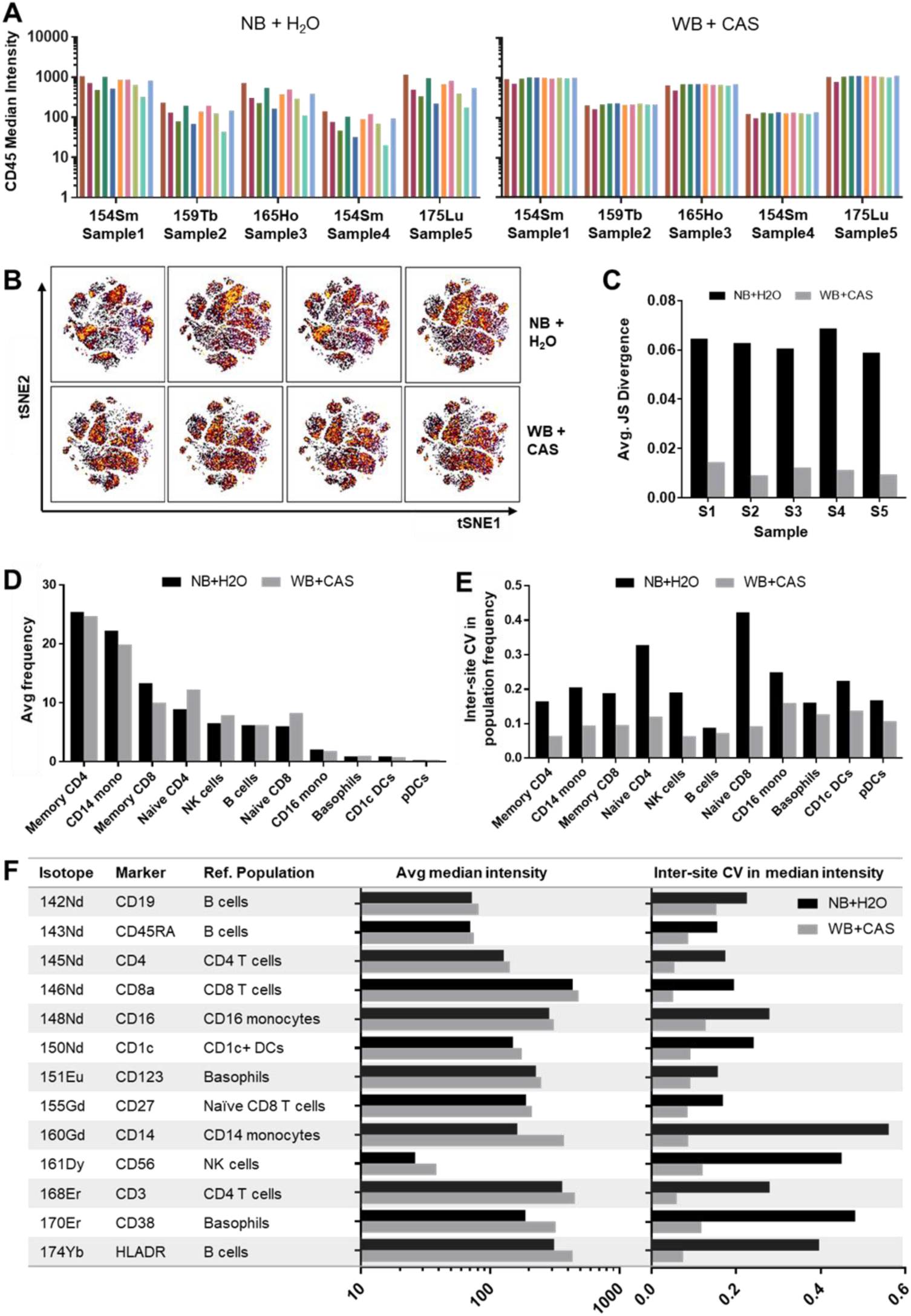
The WB+CAS protocol improves data consistency and results in an overall reduction in inter-instrument variability. (A) The individual CD45-barcoded PBMC samples as described in Fig. 5 were demultiplexed using manual Boolean gating and the median CD45 barcode intensity was determined for each sample. Each bar corresponds to a single instrument. (B) Each PBMC sample was analyzed by tSNE using all of the immunophenotyping markers in the panel to evaluate overall staining quality. Representative tSNE plots (colored by density) show data for one PBMC sample acquired on 4 different Helios instruments using either the NB+H_2_O (top) or WB+CAS (bottom) protocols. (C) Average pairwise Jensen-Shannon divergence calculated for all tSNE plots for each PBMC sample acquired using either the NB or WB protocols on all 10 instruments (p <0.01 for all samples between WB and NB). (D) Data were clustered by SPADE and major populations were identified based on canonical marker expression patterns as in Fig 2. Average frequency of indicated immune populations measured using the NB or WB protocols (averaged across all samples and instruments). (E) Inter-instrument CV in population frequency across all 10 instruments (averaged across all 5 PBMC samples; p< 0.05 for all populations between WB and NB). (F) Median marker intensity (averaged across all samples and instruments) and inter-instrument CV in marker intensity across all 10 instruments (averaged across all 5 PBMC samples; p < 0.01 for all markers).

## Discussion

The introduction of the 3^rd^ generation Helios mass cytometer offered a number of practical and theoretical advantages over the previous CyTOF2 mass cytometer, including a more efficient sample introduction system and an injector that generates smaller ion clouds, allowing higher sample throughput. However, our data highlights that this new NB injector may result in an unintended reduction in cell-associated staining quality on some Helios instruments, reflected by a reduction in median antibody signal intensity, increased variation in antibody expression within a given cell population, and reduced resolution of distinct cell subsets based on marker expression levels. We have also observed a similar phenomenon when testing the performance of the NB Helios injector installed on a CyTOF2 instrument (data not shown), further suggesting that the reduction in data quality is due to the NB injector.

The divergent behavior between the cell- and bead-derived signals detected in the same mass channel indicates that this is specifically a cell-associated phenomenon. This divergence means that these data artifacts cannot be effectively addressed by traditional bead-based normalization. Given that ion clouds derived from cells and beads would be expected to behave similarly after exiting the ICP, this suggests that the reduced cell signal intensity is due to a problem with cell stability prior to entering the plasma. While we found that multiple markers were affected across multiple cell types, we observed a trend towards a greater impact on markers at the higher end of the detectable mass range. However, we noted that the phenomenon does not appear to be specific to any particular immune cell type or due to a progressive time-dependent degradation of sample quality. Interestingly, the increase in marker staining CV suggests that some cells in a sample may be impacted to a greater degree than others, though this effect appears to be somewhat stochastic and unrelated to any specific defined cell population. Consequently, in the case of some instruments and samples, markers that may show a largely unimodal distribution within a given cell population may instead be skewed as a bimodal distribution composed of some cells that express “normal” levels and others with apparent low expression of that marker. This may result in erroneous data interpretation and false identification of cell subsets that are technical rather than truly biological in origin. This is a particular concern in the case of certain cell subsets that are traditionally defined by downregulation of a specific marker, such as “myeloid derived suppressor cells,” which are often defined as a subset of monocyte-like cells with reduced expression of HLA-DR(10).

We found that the adoption of Fluidigm’s recently-introduced wide-bore injector and CAS buffer largely mitigated these issues, resulting in generally higher signal intensities and lower signal CVs within populations, which restored data quality to levels comparable to or exceeding that found with the CyTOF2. We found that substituting the NB for WB injector alone offered some benefit to Helios data quality; however, optimal data quality was achieved when the WB injector was used in conjunction with CAS. This indicates that both the injector and acquisition buffer may independently contribute to data quality and suggests that use CAS could potentially result in improve data quality while using the NB injector. However, we did not explicitly test this condition since Fluidigm does not recommend using CAS with the NB injector due to concerns regarding the potential of residue deposition in the injector, which could have a greater impact on the narrower injector bore.

The range of inter-instrument variability that we found highlights that the NB-associated data quality artifacts are highly instrument specific; some of the instruments tested already performed well with the original NB injector protocol, and in these cases the adoption of the WB protocol had minimal effect on relative data quality. However, in cases where instruments performed poorly with the NB injector, the WB protocol mitigated this issue, resulting in the overall normalization of instrument performance and improving the consistency of data between instruments. Our results suggest that the WB injector may also potentially impact the reproducibility of data acquired on a given instrument over time, but this remains to be comprehensively tested. When evaluating instrument tuning and performance parameters that may be associated with these data quality issues, we observed a positive correlation between the relative impact of the WB injector protocol and Tb(159) dual counts at tuning, suggesting that instruments that are ostensibly showing greater sensitivity may in fact be more prone to data artifacts when using the NB injector, and would potentially benefit most from adoption of the WB protocol. While the adoption of the WB protocol has the potential to improve data quality from poorly performing instruments, this improvement comes at the expense of throughput, since the WB injector results in a higher relative doublet rate, which necessitates a lower acquisition rate to ensure maximal singlet recovery. The ionic content of the CAS buffer also results in increased residue deposits on the injector during normal operation. While this can be easily addressed by daily cleaning of the injector, this does represent an additional burden of instrument maintenance.

In addition to being instrument specific, our data suggest that the data artifacts associated with the NB injector and the consequent relative improvement in performance with the WB protocol modification may also vary on the same instrument over time and may also be somewhat sample specific (data not shown). The sample aliquots used in our multi-site comparison studies were stained, fixed and then cryopreserved using FBS+10% DMSO prior to distribution to each site in order to allow for a more streamlined and standardized sample acquisition protocol and minimize other potential sources of technical variation. While we and others have previously shown that this is an effective method to preserve stained samples prior to analysis using the CyTOF2(2,3), we present evidence that cryopreserved samples may be particularly sensitive to the data quality issues associated with the NB protocol on some Helios instruments but not others.

Overall, these results clearly demonstrate the potential value of the WB injector protocol in improving data quality on some Helios instruments. This is particularly valuable in the case of multi-site studies, where we found that adoption of the WB protocol ultimately resulted in high levels of inter-instrument reproducibility, with inter-instrument CVs <10% for most cell population frequencies and individual marker expression levels within those populations when measuring parallel aliquots of the same pre-stained sample. However, these results also highlight that both the data artifacts associated with the NB injector and the relative performance improvements seen with the WB injector are both somewhat instrument and sample specific. While adoption of WB injector comes at the cost of reduced throughput and increased maintenance requirements, our findings strongly support that any site currently performing mass cytometry studies with a Helios mass cytometer would benefit from evaluating the impact of the two protocols on their specific instrument. This is particularly important in the case of multi-center studies where data are being collected across multiple instruments. Our findings also highlight the importance of using relevant biological standards both to monitor inter-instrument variability in multi-center studies, and to rigorously test and benchmark the consequences of any instrument modifications or upgrades.

## Acknowledgements

We thank Seunghee Kim-Schulze and the Mt. Sinai Human Immune Monitoring Center for technical support. We also thank Mike Leipold, Natalia Sigal and Holden T. Maecker for their contributions to the multi-site testing. This work includes data collected on a Helios mass cytometry instrument supported by S10 OD023547-01. The multicenter comparison studies include data generated through the NCI Cancer Immune Monitoring and Analysis Centers (CIMAC) CyTOF Assay Working Group and the Human Immunology Project Consortium (HIPC) Samples and Assays Subcommittee and were partially supported by U19AI118610, U19AI128949, U19AI118626 and U24CA224319. Fluidigm provided WB injectors and CAS buffer for initial testing and evaluation.

