## Supplementary materials for "A Modified Injector and Sample Acquisition Protocol Can Improve Data Quality and Reduce Inter-Instrument Variability of the Helios Mass Cytometer"

**Supplementary Table 1. Antibodies**

| Immunophenotyping antibodies | | | |
|---|---|---|---|
| **Isotope** | **Target** | **Clone** | **Source** |
| 142 Nd | CD19 | HIB19 | Biolegend |
| 143 Nd | CD45RA | HI100 | Biolegend |
| 145 Nd | CD4 | RPA-T4 | Biolegend |
| 146 Nd | CD8a | RPA-T8 | Biolegend |
| 148 Nd | CD16 | 3G8 | Biolegend |
| 150 Nd | CD1c | L161 | Biolegend |
| 151 Eu | CD123 | 6H6 | Biolegend |
| 152 Sm | CD66b | G10F5 | Biolegend |
| 155 Gd | CD27 | O323 | Biolegend |
| 160 Gd | CD14 | M5E2 | Biolegend |
| 161 Dy | CD56 | B159 | BD Biosciences |
| 164 Dy | CD161 | HP-3G10 | Fluidigm |
| 165 Ho | CD45 | HI30 | Biolegend |
| 168 Er | CD3 | UCHT1 | Biolegend |
| 169 Tm | CD45 | HI30 | Biolegend |
| 170 Er | CD38 | HB-7 | Biolegend |
| 172 Yb | CD57 | HCD57 | Fluidigm |
| 174 Yb | HLADR | L243 | Biolegend |
| 175 Lu | CD45 | HI30 | Biolegend |
| 176 Yb | CD56 | NCAM16.2 | Fluidigm |
| 209 Bi | CD11b | ICRF44 | Fluidigm |
| **CD45 Barcoding antibodies** | | | |
| **Isotope** | **Target** | **Clone** | **Source** |
| 89 Y | CD45 | HI30 | Biolegend |
| 113 In | CD45 | HI30 | Biolegend |
| 115 In | CD45 | HI30 | Biolegend |
| 141 Pr | CD45 | HI30 | Biolegend |
| 154 Sm | CD45 | HI30 | Biolegend |
| 159 Tb | CD45 | HI30 | Biolegend |
| 194 Pt | CD45 | HI30 | Biolegend |
| 198 Pt | CD45 | HI30 | Biolegend |

**Supplementary Table 2. Tuning parameters and their association with the relative change in data quality between the NB and WB protocols**

| Helios Instrument | FC in Median (WB/NB) | FC in CV (WB/NB) | Nebulizer gas (L/min) | | | Makeup gas (L/min) | | | Tb(159) Mean Dual Counts | | | Tb(159) Mean RSD | | | Gd(155)/La(139) oxide ratio | | |
|---|---|---|---|---|---|---|---|---|---|---|---|---|---|---|---|---|---|
| | | | NB | WB | WB/NB ratio | NB | WB | WB/NB ratio | NB | WB | WB/NB ratio | NB | WB | WB/NB ratio | NB | WB | WB/NB ratio |
| A | -0.124 | -0.008 | 0.17 | 0.18 | 1.06 | 0.49 | 0.74 | 1.51 | 634078 | 897168 | 1.41 | 1.024 | 0.498 | 0.49 | 0.02 | 0.021 | 1.02 |
| B | 0.345 | -0.103 | 0.25 | 0.25 | 1.00 | 0.42 | 0.66 | 1.57 | 1364156 | 1501782 | 1.1 | 0.715 | 0.528 | 0.74 | 0.035 | 0.046 | 1.3 |
| C | 0.887 | -0.204 | 0.16 | 0.15 | 0.94 | 0.51 | 0.79 | 1.55 | 1682876 | 1551329 | 0.92 | 0.287 | 0.306 | 1.07 | 0.061 | 0.043 | 0.71 |
| D | 2.225 | -0.264 | 0.17 | 0.14 | 0.82 | 0.48 | 0.68 | 1.42 | 1675695 | 2026936 | 1.21 | 0.419 | 0.385 | 0.92 | 0.049 | 0.06 | 1.22 |
| E | 0.147 | -0.117 | 0.16 | 0.16 | 1.00 | 0.47 | 0.69 | 1.47 | 1114502 | 1135357 | 1.02 | 0.499 | 0.373 | 0.75 | 0.011 | 0.011 | 1.03 |
| F | 0.117 | -0.043 | 0.26 | 0.26 | 1.00 | 0.4 | 0.6 | 1.5 | 1266890 | 1194119 | 0.94 | 0.76 | 0.516 | 0.68 | 0.034 | 0.025 | 0.74 |
| G | 0.656 | -0.142 | 0.18 | 0.17 | 0.94 | 0.47 | 0.69 | 1.47 | 1143005 | 1043363 | 0.91 | 0.803 | 0.585 | 0.73 | 0.064 | 0.05 | 0.79 |
| H | 0.249 | -0.106 | 0.15 | 0.15 | 1.00 | 0.47 | 0.67 | 1.43 | 955780 | 807415 | 0.84 | 0.165 | 0.431 | 2.6 | 0.045 | 0.04 | 0.89 |
| I | 1.391 | -0.194 | 0.16 | 0.16 | 1.00 | 0.54 | 0.77 | 1.43 | 1128532 | 1517862 | 1.34 | 0.439 | 0.665 | 1.52 | 0.02 | 0.025 | 1.27 |
| J | 0.073 | -0.099 | 0.17 | 0.16 | 0.94 | 0.48 | 0.73 | 1.52 | 747082 | 888147 | 1.19 | 0.612 | 0.331 | 0.54 | 0.054 | 0.045 | 0.83 |
| **Correlation with FC in median** | | | -0.24 | -0.42 | -0.77 | 0.42 | 0.21 | -0.50 | 0.68 | 0.86 | 0.19 | -0.42 | 0.06 | 0.16 | 0.24 | 0.52 | 0.43 |
| **Correlation with FC in CV** | | | 0.39 | 0.56 | 0.79 | -0.49 | -0.34 | 0.39 | -0.73 | -0.78 | 0.04 | 0.63 | 0.15 | -0.23 | -0.37 | -0.54 | -0.28 |

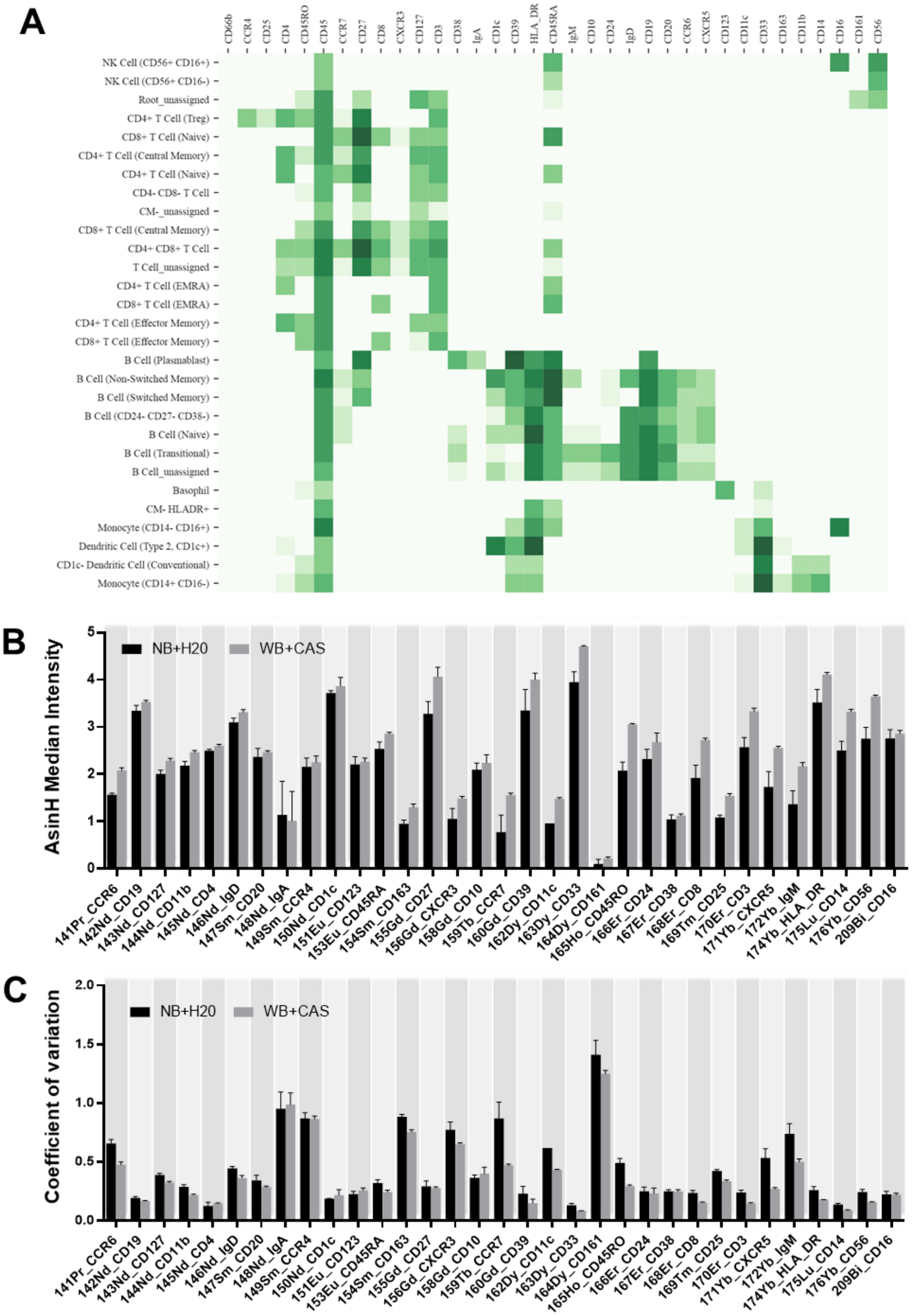


**Supplementary Figure 1. Independent validation of the impact of the NB+$H_2O$ and WB+CAS protocols on Helios data quality**. PBMC samples from three donors were stained and acquired as three independent technical replicates using either the NB or WB protocols. The samples were processed and acquired at an independent site using a different Helios mass cytometer from those used to generate the data shown in Figs.1-4. The resulting data were clustered and annotated using flowSOM as implemented in Astrolabe. (A) Representative Astrolabe heatmap of identified population clusters showing median marker expression across clusters. Median marker intensity (B) and marker CV (C) as determined for the highest-expressing population cluster for each marker. Bars represent the mean + S.D. for the three samples run in triplicate on one instrument under each injector protocol.

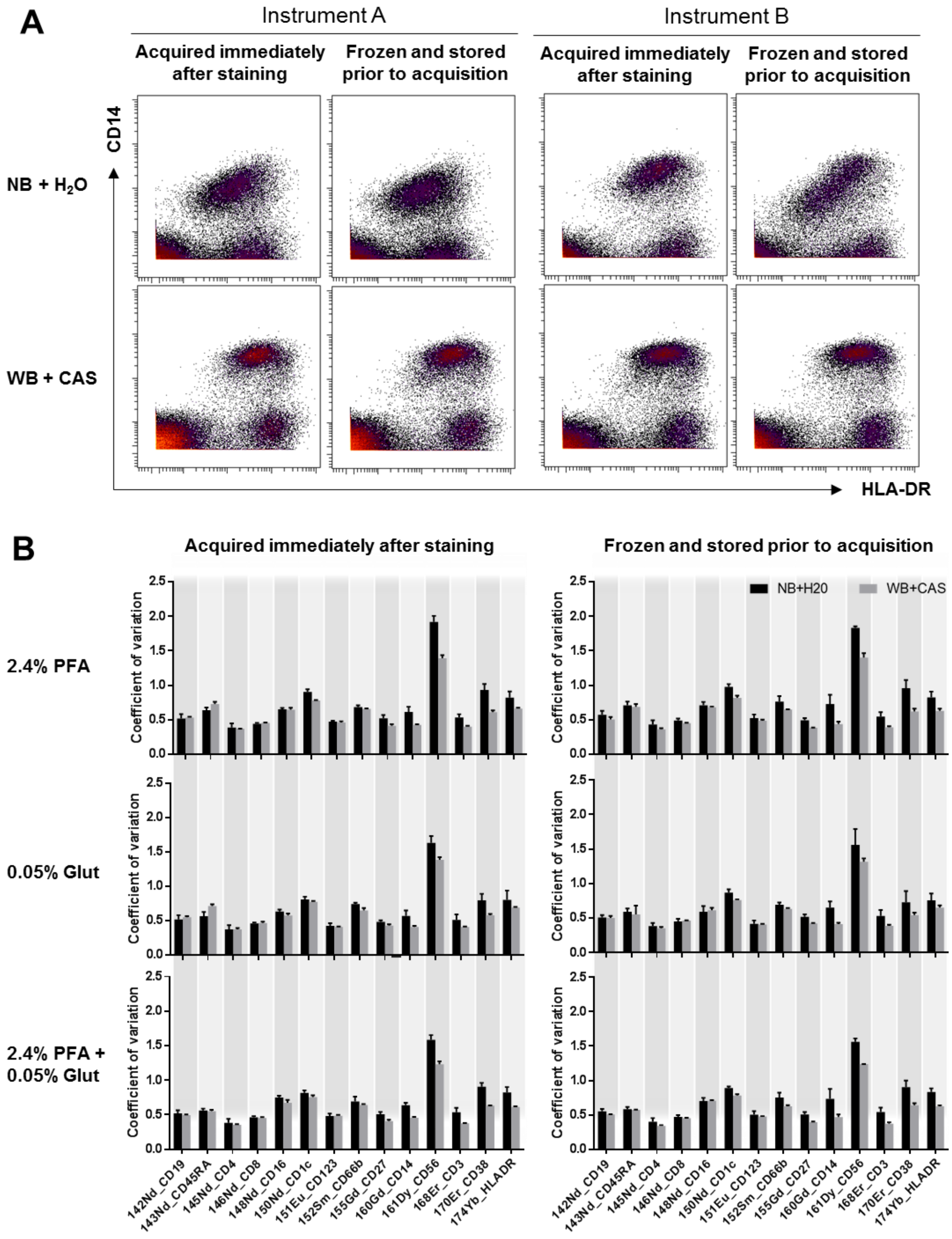


**Supplementary Figure 2. The WB+CAS protocol improves Helios data quality under a range of sample fixation and storage conditions.** PBMCs from a single donor were stained with an antibody panel and then fixed for 30 minutes using 2.4% formaldehyde, 0.05% glutaraldehyde or a combination of 2.4% formaldehyde and 0.0% glutaraldehyde in PBS. Replicate aliquots of the stained and fixed sample were acquired on three independent Helios instruments either immediately following staining or after cryopreservation and storage at -80oC in 10% DMSO in FBS. (A) Representative biaxial plots showing data from formaldehyde-fixed samples that were acquired either immediately after staining or after cryopreservation and storage, on two different instruments using either the NB or WB protocols. The data illustrate further exacerbation of the NB-associated data quality issues with the cryopreserved samples on one instrument (right), but not the other (left); however, the WB+CAS protocol shows improvement in both cases. (B) The data were analyzed by SPADE using the same strategy and reference populations as described in figure 2. The graphs show the coefficient of variation of each marker on its designated reference population using the NB+H2O or WB+CAS protocols (bars represent mean + SD of triplicate aliquots acquired on three instruments).

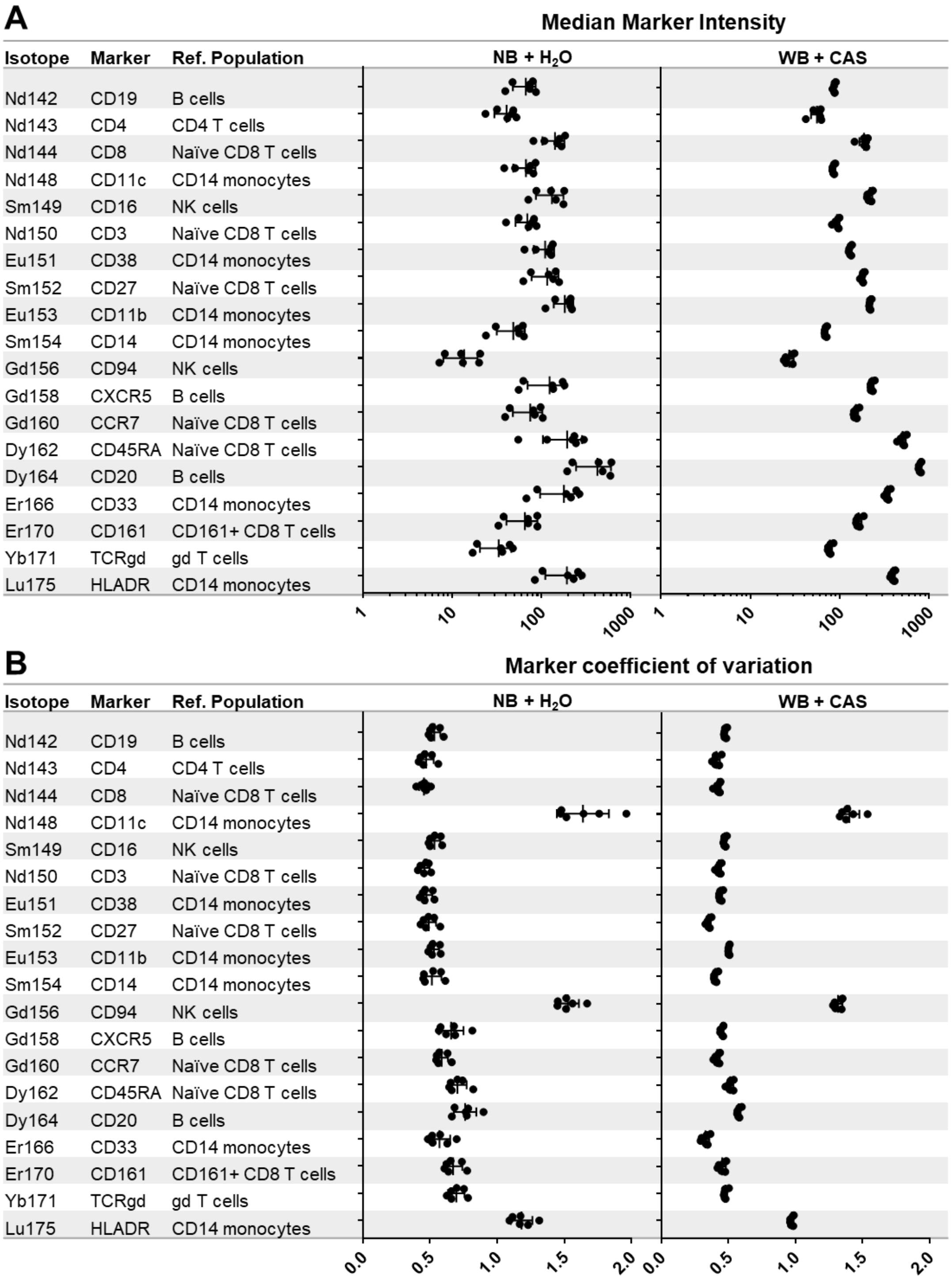


**Supplementary Figure 3. Secondary validation of the impact of the NB+$H_2O$ and WB+CAS protocols on inter-instrument data reproducibility.** An independent PBMC samples was stained and processed at a second site using a distinct antibody panel from that used in the primary multi-site comparison study, and frozen aliquots of the pre-stained cells were distributed and acquired on 6 Helios instruments (a subset of those used in the primary multi-site comparison) using either the NB+$H_2O$ or the WB+CAS acquisition protocols. Major immune populations were manually gated, and a positive reference population was designated for each marker. Graphs represent the median marker intensity (A) and coefficient of variation (B) for each of the indicated marker-population combinations. Each point represents a separate instrument, highlighting the overall reduction in inter-instrument variation with the WB+CAS protocol.
